# PIEZO1 Drives Trophoblast Fusion and Placental Development

**DOI:** 10.1101/2025.03.25.645313

**Authors:** Yang Zhang, Ke Z. Shan, Pengfei Liang, Augustus J. Lowry, Liping Feng, Huanghe Yang

## Abstract

PIEZO1, a mechanosensor^1,2^ in endothelial cells, plays a critical role in fetal vascular development during embryogenesis^3,4^. However, its expression and function in placental trophoblasts remain unexplored. Here, we demonstrate that PIEZO1 is expressed in placental villus trophoblasts, where it is essential for trophoblast fusion and placental development. Mice with trophoblast-specific PIEZO1 knockout exhibit embryonic lethality without obvious vascular defects. Instead, PIEZO1 deficiency disrupts the formation of the syncytiotrophoblast layer in the placenta. Mechanistically, PIEZO1-mediated calcium influx activates TMEM16F lipid scramblase, facilitating the externalization of phosphatidylserine, a key “fuse-me” signal for trophoblast fusion^5,6^. These findings reveal PIEZO1 as a crucial mechanosensor in trophoblasts and highlight its indispensable role in trophoblast fusion and placental development, expanding our understanding of PIEZO1’s functions beyond endothelial cells during pregnancy.

## Main Text

Mechanical forces are essential regulators of embryonic development, shaping processes from gastrulation to organogenesis ^7–10^. In contrast, mechanobiology of the placenta, a vital yet understudied organ ^11^, remains poorly defined. During pregnancy, the placenta undergoes profound mechanical adaptations, including trophoblast migration, decidual invasion, syncytialization (trophoblast fusion), villous morphogenesis, and spiral artery remodeling ^7,12–14^. While mechanical cues drive these processes, the molecular sensors transducing forces into biochemical signals in trophoblasts remain elusive.

PIEZO1, a mechanosensitive ion channel ^1,2^, has emerged as a key force transducer in development. Constitutive PIEZO1 knockout (KO) mice are embryonically lethal, exhibiting severe defects in fetal angiogenesis ^3,4^. Similarly, endothelial-specific deletion of PIEZO1 recapitulates this phenotype, underscoring its essential role in endothelial mechanotransduction. Beyond vascular development, PIEZO1 also regulates neural stem cell lineage specification ^15–17^, osteogenesis ^18,19^, and lymphangiogenesis ^20,21^. However, despite trophoblasts being exposed to dynamic shear stress and/or compressive forces during gestation ^14^, whether PIEZO1 mediates force sensing in trophoblasts remains unknown. Here, we identify PIEZO1 as a mechanosensor in villous trophoblasts and demonstrate its essential role in trophoblast fusion and placental development. By shifting the focus from endothelial cells to trophoblasts, this study reveals a previously unrecognized PIEZO1-dependent mechanotransduction pathway in trophoblasts, providing insights into the mechanical regulation of the placenta development and identifying potential therapeutic targets for pregnancy complications associated with placenta dysfunction. **Results**

### PIEZO1 is expressed in placental trophoblasts

Immunofluorescence analysis revealed that PIEZO1 is expressed in both single-nucleated cytotrophoblasts (CTB) and multi-nucleated syncytiotrophoblasts (STB) in human chorionic villi (Fig. 1A), as well as in the human placental trophoblast cell line, BeWo (Fig. 1B). Pressure clamp electrophysiology recording elicited a mechanosensitive current from BeWo cells with a unitary conductance of 29.3 ± 1.8 pS (Fig. 1C-G), which aligns with the typical single channel conductance of PIEZO1 ^2,22^. Yoda1, a PIEZO1 agonist, robustly increased intracellular Ca^2+^ ([Ca^2+^]_i_), which can be attenuated by the PIEZO1 inhibitor, GsMTx4 (Fig. S1). Silencing PIEZO1 using siRNA not only markedly attenuated this mechanosensitive current (Fig. 1E-G) but also reduced Yoda1-induced [Ca^2+^]_i_ increase (Fig. 1H-I). Taken together, this evidence supports that PIEZO1 is functionally expressed in human trophoblasts.

**Fig. 1.**
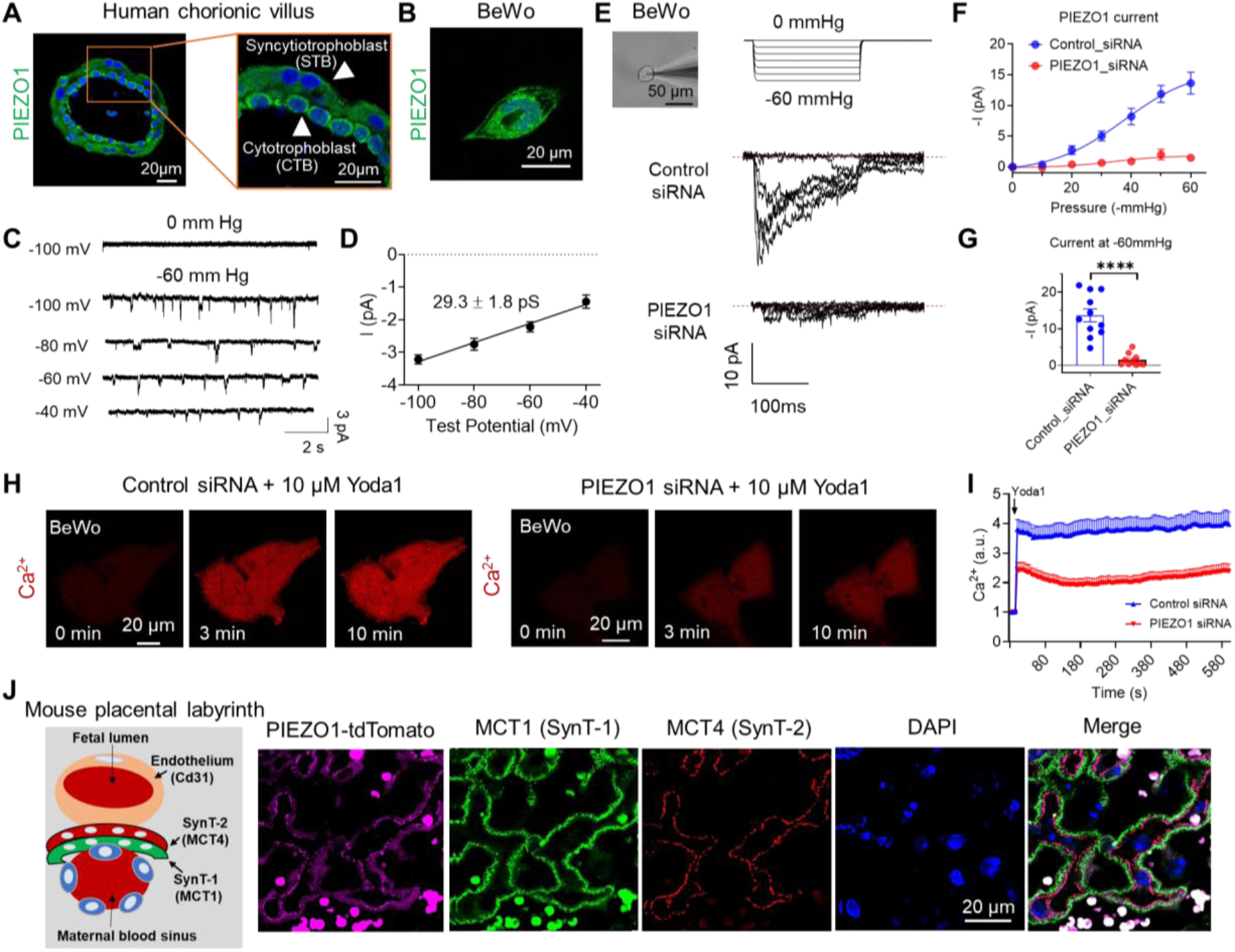
PIEZO1 is expressed in placental trophoblasts. **(A)** Immunofluorescence staining of PIEZO1 (green) and nuclei (Hoechst, blue) in human first-trimester placenta villi with higher magnification shown on the right. **(B)** Immunofluorescence staining of PIEZO1 (green) in BeWo cells. **(C)** Representative single channel mechanosensitive currents from BeWo cells. Currents were elicited by -60 mmHg pressure and recorded at voltage steps from -100 mV to -40 mV in 20 mV increments with holding potential at 0 mV. **(D)** The current-voltage (I-V) relationship for the currents in (C). The data were fitted with linear regression, obtaining a single-channel conductance of 29.3 ± 1.8 pS (n = 5). **(E)** Representative cell-attached pressure-clamp recording of control-siRNA and PIEZO1-siRNA treated BeWo cells (left). The macroscopic current was elicited by pressure steps ranging from 0 to −60 mmHg with a holding potential at −80 mV. **(F)** The current-pressure relationship for the mechanosensitive current recorded from BeWo cells treated with control-siRNA and PIEZO1-siRNA. **(G)** PIEZO1 current amplitudes at –60 mmHg in (F). Unpaired 2-sided Student t-test. ∗∗∗∗: P < 0.0001, n = 10-11 for each condition. **(H)** Representative Ca^2+^ (red) imaging of control and PIEZO1 siRNA knockdown BeWo cells stimulated with 10 µM Yoda1. **(I)** Time course quantification of Ca^2+^ levels in control (n=21) and PIEZO1 siRNA (n=25) knockdown BeWo cells following 10 µM Yoda1 stimulation (indicated by the arrow). Data are presented as mean ± s.e.m from 7 biological replicates. **(J)** Immunofluorescence of tdTomato (magenta), MCT1 (a marker for SynT-1 layer, green), MCT4 (a marker for SynT-2 layer, red), and DAPI (blue) for the placenta from Piezo1-td-Tomato transgenic mouse at E13.0. Left: schematic of mouse placental labyrinth showing the maternal-fetal interface. Circular tdTomato+ and MCT1+ signals are red blood cells. n = 4 biological replicates.

We previously reported that the TRPV4 ion channel is specifically expressed in human but not mouse trophoblasts, indicating differential expression profiles in trophoblasts from different species ^23^. To determine whether Piezo1 is also expressed in mouse trophoblasts, we examined its protein expression in the mouse placenta utilizing the Piezo1-tdTomato reporter mice (Piezo1^P1-tdT^) in which tdTomato is covalently tagged to endogenous Piezo1 ^4^. The td-Tomato signal was detected in the placental labyrinth where SynT-1 and SynT-2, two layers of STBs, are located (Fig. 1J) ^24^. We found strong td-Tomato signal in SynT-2 and weaker signal in SynT-1 using antibodies for MCT1 and 4, which are two monocarboxylate transporters specific to SynT-1 and -2 ^5,23,25^. Consistently, single-cell RNA sequencing of the mouse placenta from the Spatiotemporal Transcriptomic Atlas of Mouse Placenta (STAMP) ^26^ also shows that *Piezo1* (Fig. S2A-B) is highly expressed in SynT-2 STBs that specifically express the fusogenic gene *Synb*, but weakly expressed in SynT-1 STBs that specifically express another fusogenic gene *Syna* (Fig. S2C-D) ^24^. Taken together, our immunostaining and functional analysis confirm that, unlike TRPV4, PIEZO1 is expressed in both human and mouse placental trophoblasts and predominantly localizes to SynT-2 STBs in the mouse labyrinth.

### Trophoblast-specific deletion of Piezo1 results in embryonic lethality

Piezo1 constitutive KO mice are embryonically lethal, a phenotype previously attributed to endothelial dysfunction and defective angiogenesis due to PIEZO1 deficiency ^3,4^. To dissect the role of PIEZO1 in placental trophoblasts, we generated a trophoblast-specific knockout (cKO) of Piezo1 mice by breeding *Piezo1^f/f^*^27^ with *Elf5*-*Cre,* a transgenic mouse line that specifically drives Cre recombinase expression in the trophectoderm as early as embryonic day 4.5 (E4.5) ^28^. As illustrated in the breeding scheme in Fig. 2A, the anticipated Mendelian ratio to obtain cKO is 25% (Fig. 2B, left). Nevertheless, nearly all cKO embryos died during pregnancy, with only 1 out of 105 (<1%) cKO mice surviving after birth (Fig. 2B, right). The cKO embryos and placentas were markedly smaller compared to their litter mates; some even showed signs of degradation and resorption (Figs. 2C and S3), suggesting that cKO mice suffer from intrauterine growth restriction (IUGR) and stillbirth. Embryonic death of Piezo1 trophoblast cKO mice starts as early as E11.5 (Fig. S3A).

**Fig. 2.**
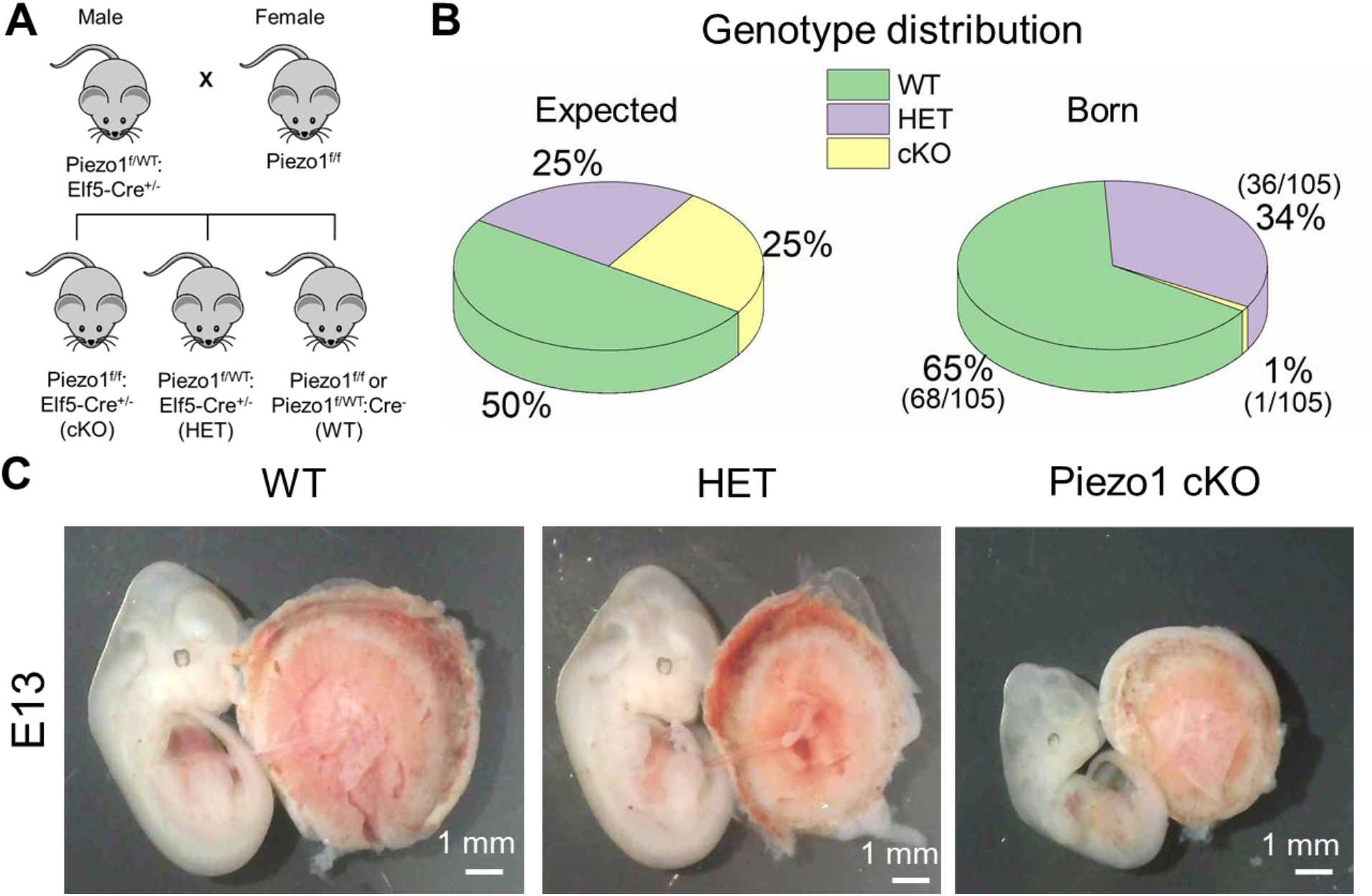
Trophoblast-specific knockout of Piezo1 causes embryonic lethality in mice. **(A)** Breeding scheme for the generation of Piezo1 conditional knockout (cKO) mice. **(B)** Genotype distribution of wild-type (WT), heterozygous (HET), and cKO pups compared to the expected Mendelian inheritance ratio. The numbers of animals with specific genotypes over 105 pups were shown in the parentheses. **(C)** Representative embryos and placentas from Piezo1 WT, HET, and cKO mice at embryonic day 13.0 (E13.0). Images of the complete litter can be found in Fig. S3B.

### Piezo1 deficiency disrupts trophoblast fusion

We conducted histological analysis to identify the defects in the cKO placenta. H&E staining demonstrated that E13.0 cKO placentas had markedly enlarged maternal blood sinuses in the labyrinth compared to those in WT placentas from the same litter (Fig. S4, red asterisks). Interestingly, the numbers of fetal capillaries, which were identified by nucleated fetal red blood cells (RBCs), did not show an obvious difference between WT and cKO placentas (Fig. S4, blue stars). Our CD31 staining of WT and cKO placentas further supports that trophoblast-specific deletion of Piezo1 does not significantly impact fetoplacental blood vessel development (Fig. 3A-B). This is distinct from the severe angiogenesis defects observed in the constitutive Piezo1 KO and the endothelial-specific Piezo1 cKO placentas ^3,4^. Instead, our histological analysis suggests that defective trophoblast differentiation, rather than impaired angiogenesis, is the primary cause of embryonic lethality in trophoblast-specific Piezo1 cKO mice.

**Fig. 3.**
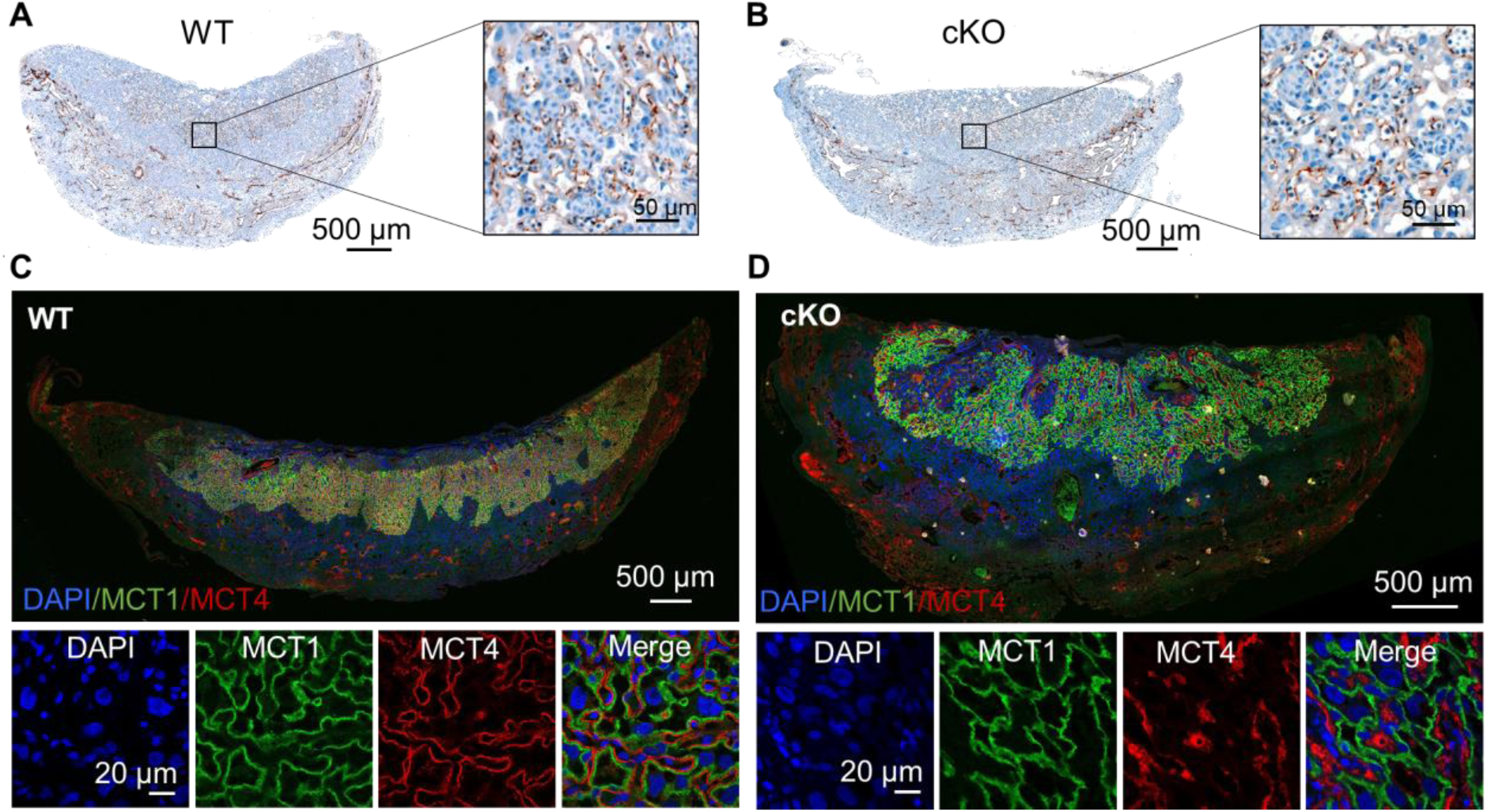
Piezo1 deficiency in trophoblast impairs fusion without affecting angiogenesis. **(A-B)** CD31 immunohistochemical staining of PIEZO1 WT (A) and cKO (B) placentas at E13.0 with enlarged areas shown on the right. n = 3 biological replicates for WT and 3 biological replicates for cKO. **(C-D)** MCT1 and MCT4 immunofluorescence staining of the PIEZO1 WT (C) and cKO (D) placentas (upper panels) with enlargement (lower panels) at E12.5. n = 5 biological replicates for WT and 6 biological replicates for cKO.

Defects in trophoblast fusion, a crucial step of trophoblast differentiation, are a major contributor to placental insufficiency, IUGR, and in severe cases, stillbirth ^29–31^. To evaluate STB formation in the cKO labyrinth, we stained MCT1 and MCT4, markers of the SynT-1 and SynT-2 STBs, respectively ^5,23,25^. In stark contrast to the smooth, uniform, and colocalized MCT1 and MCT4 staining in WT labyrinth, MCT4 expression was dramatically reduced, and in some regions, entirely absent in the Piezo1 cKO labyrinth. Although MCT1 staining was largely preserved, it exhibited a more irregular, zigzag pattern (Fig. 3C-D). These findings indicate that trophoblast-specific deletion of Piezo1 severely disrupts the formation of the SynT-2 STB layer while having a milder effect on the SynT-1 STB layer, consistent with Piezo1’s expression profile in these STB layers (Figs. 1J and S2).

### PIEZO1 is essential for trophoblast fusion *in vitro*

To further confirm the essential role of PIEZO1 in trophoblast fusion, we pharmacologically and genetically manipulated PIEZO1 in BeWo cells and quantified its effects on forskolin-induced BeWo cell fusion *in vitro*. Inhibition of PIEZO1 with GsMTx4 abolished BeWo cell fusion (Fig. 4A-B). Similarly, PIEZO1 knockdown via siRNA significantly reduced the fusion index (Fig. 4C-D). Conversely, PIEZO1 overexpression significantly increased PIEZO1 current (Fig. S5) and enhanced BeWo cell fusion index from 0.42±0.03 to 0.65±0.02 (Fig. 4E-F). Our *in vitro* experiments in a human trophoblast cell line thus support our *in vivo* findings in the mouse placenta, confirming that PIEZO1 is indispensable for trophoblast fusion.

**Fig. 4.**
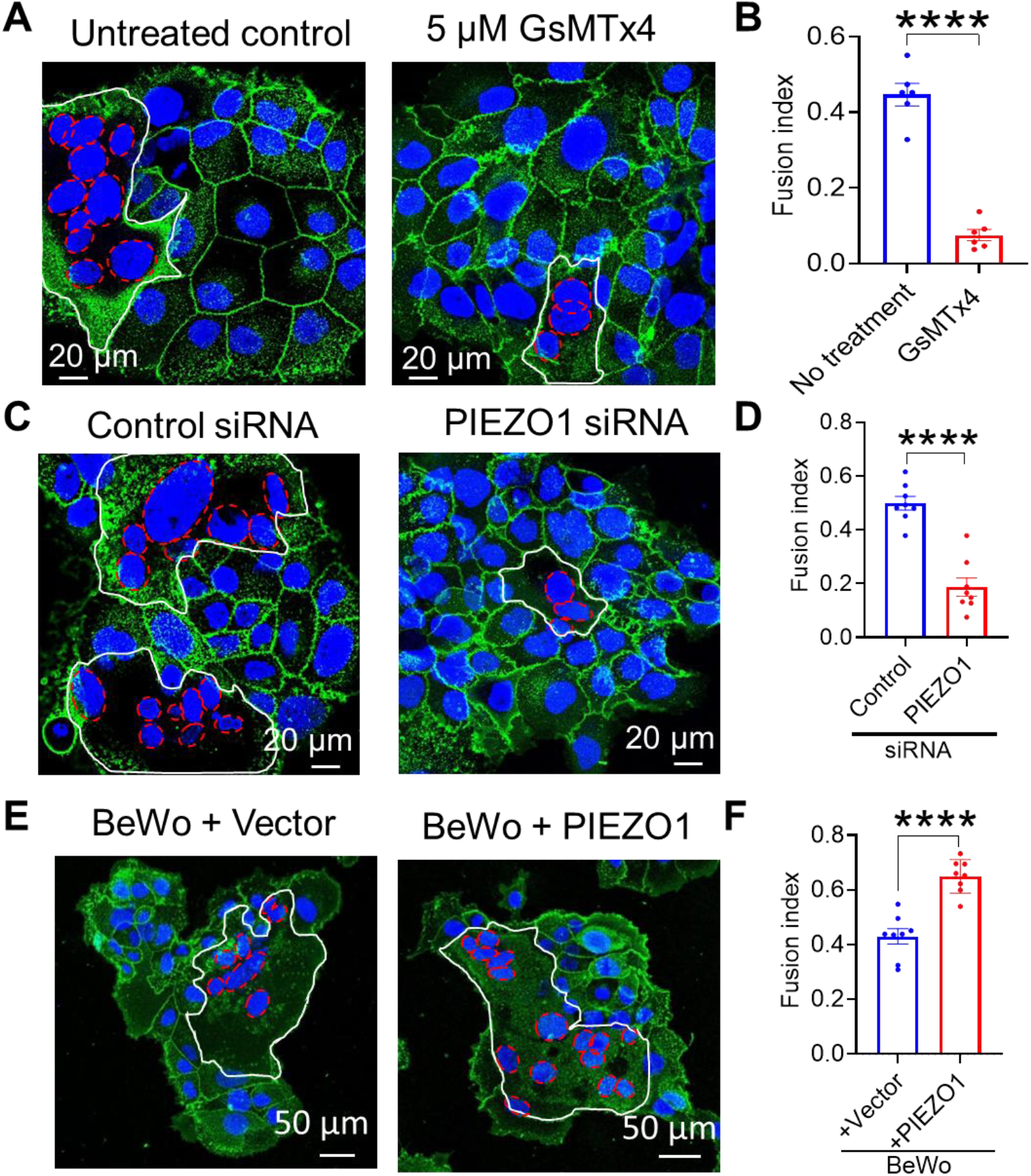
PIEZO1 is required for trophoblast fusion in vitro. **(A-B)** Representative images (A) and fusion index quantification (B) of untreated control (n=6) and 5 μM GsMTx4 treated (n=6) BeWo cells after 48-hr forskolin treatment. **(C-D)** Representative images (C) and fusion index quantification (D) of control (n=8) and PIEZO1 siRNA knockdown (n=8) BeWo cells after forskolin treatment for 48-hr to induce fusion. **(E-F)** Representative images (E) and fusion index quantification (F) of BeWo cells that were overexpressed with empty vector (n=8) and PIEZO1 (n=8) after 48-hr forskolin treatment. Each dot represents fusion indexes averaged from six random fields of one coverslip. Data are presented as mean ± s.e.m. ****P<0.0001 (unpaired two-tailed t-test).

### PIEZO1-TMEM16F coupling regulates trophoblast fusion

We previously reported that TMEM16F, a Ca^2+^-activated phospholipid scramblase (CaPLSase), plays a critical role in trophoblast fusion and placental development by mediating the externalization of phosphatidylserine (PS), an essential ‘fuse-me’ signal for cell fusion ^5,6^. Subsequently, we demonstrated that Ca^2+^ influx through Ca^2+^ permeable channels such as TRPV4 can activate TMEM16F, and the functional coupling between TRPV4 and TMEM16F modulates trophoblast fusion ^23^. In our recent investigations of RBCs, we found that PIEZO1-mediated Ca^2+^ influx activates TMEM16F and enhanced PIEZO1-TMEM16F coupling contributes to the pathophysiology of RBC disorders such as hereditary xerocytosis ^32^ and sickle cell disease (see accompanied manuscript that is currently under review in *Blood*). Interestingly, *Piezo1* and *Tmem16f* transcripts are both highly expressed in the SynT-2 layer (Fig. S2B-E), and the deficiency of either protein results in embryonic or perinatal lethality ^5^ accompanied by severe defects in SynT-2 STB layer development (Figs. 2 and 3C-D). Based on these findings, we hypothesized that PIEZO1 and TMEM16F could be functionally coupled in trophoblasts and this coupling plays a key role in trophoblast fusion by regulating PS exposure.

To test this hypothesis, we employed a fluorescence imaging-based CaPLSase assay ^33^ to investigate PIEZO1-TMEM16F coupling in BeWo cells. Treatment with 10 μM Yoda1 induced robust Ca²⁺ influx in both TMEM16F WT and KO BeWo cells (Fig. 5A). However, Yoda1-induced PS exposure was observed only in WT but not KO cells (Fig. 5A-B), indicating that Ca^2+^ influx through PIEZO1 can mediate TMEM16F-dependent PS exposure. On the other hand, gene silencing or GsMTx4 inhibition of PIEZO1 abolished both Yoda1-induced Ca^2+^ elevation and PS exposure in BeWo cells (Figs. 5C-D and S6). These findings demonstrate that PIEZO1 and TMEM16F are functionally coupled to regulate PS exposure in trophoblasts (Fig. 5G).

**Fig. 5.**
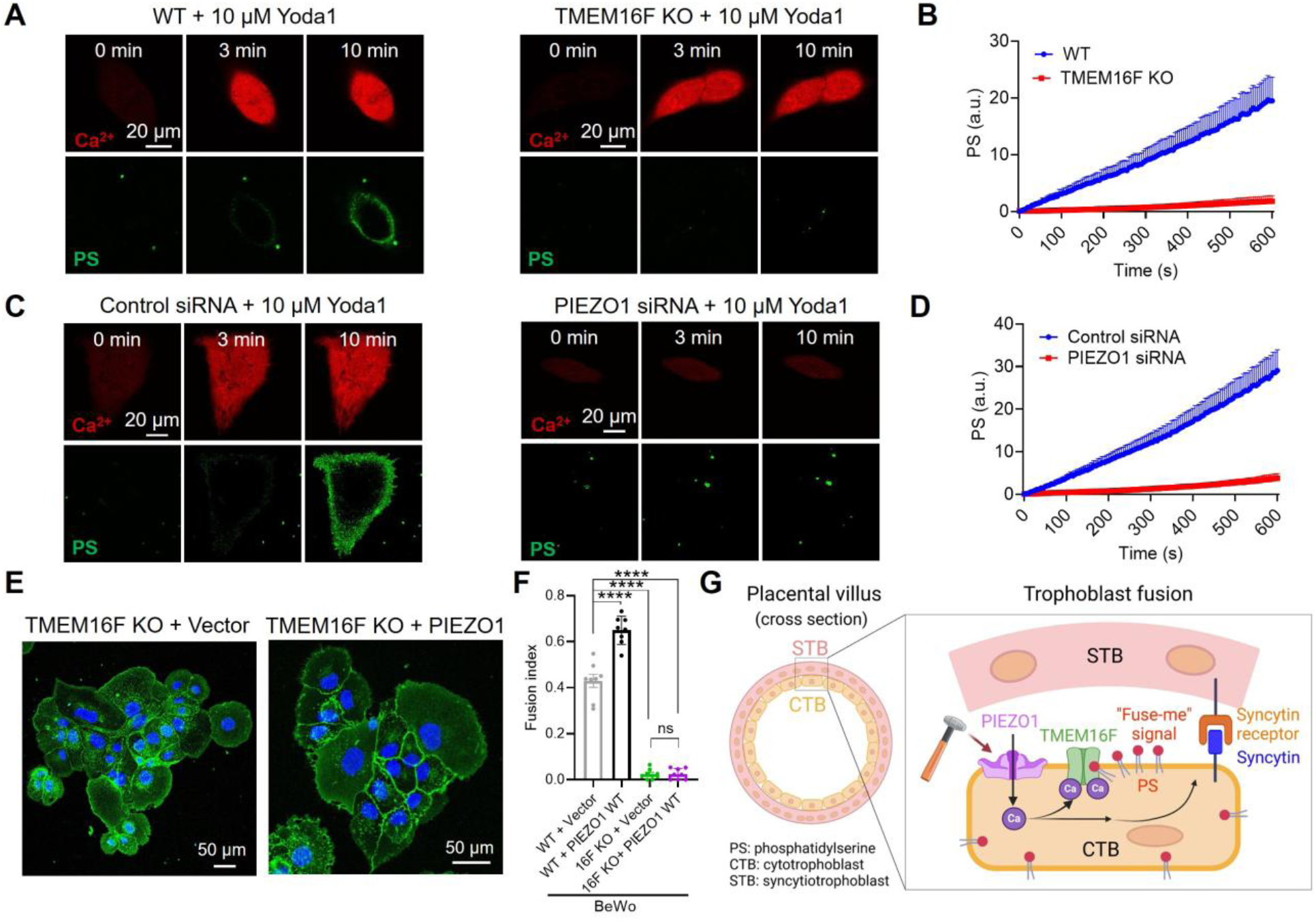
PIEZO1-TMEM16F coupling regulates trophoblast fusion. **(A)** Representative images of Ca^2+^ and PS exposure in WT (left) and TMEM16F KO (right) BeWo cells stimulated with 10 μM Yoda1. **(B)** Time course quantifications of Yoda1-induced PS exposure in WT (n= 25) and TMEM16F KO (n=18) BeWo cells. **(C)** Representative images of Ca^2+^ and PS exposure in control (left) and PIEZO1 siRNA knockdown (right) BeWo cells stimulated with 10 μM Yoda1. **(D)** Time course quantifications of Yoda1-induced PS exposure control (n= 21) and PIEZO1 siRNA (n=25) knockdown BeWo cells. **(E)** Representative images of TMEM16F KO BeWo overexpressed with empty vector and PIEZO1 after forskolin treatment for 48 hrs. **(F)** Fusion index quantification in WT and TMEM16F KO BeWo cells, which were overexpressed with an empty vector or PIEZO1 (n=8 each) after forskolin treatment for 48 hrs. Data are presented as mean ± s.e.m. n.s.; non-significant; ****P<0.0001 (one-way ANOVA with Tukey’s multiple comparisons test). **(G)** Cartoon illustration of PIEZO1-TMEM16F coupling in regulating trophoblast fusion.

As Yoda1 is not a physiological activator of PIEZO1, we next examined whether mechanical forces could directly trigger PIEZO1-TMEM16F coupling in BeWo cells. We leveraged TMEM16F’s moonlighting function as a Ca^2+^-activated ion channel ^34^ and applied a - 50 mmHg suction to a cell-attached membrane patch using a pressure clamp device. This mechanical stimulation elicited a time- and voltage-dependent, outward-rectifying current, a biophysical hallmark of TMEM16F current (Fig. S7A-B). Notably, this force-induced TMEM16F current depends on extracellular Ca^2+^ and requires the expression of both PIEZO1 and TMEM16F in BeWo cells (Fig. S7C), providing direct evidence that mechanical forces can directly activate PIEZO1-TMEM16F coupling in trophoblasts (Fig. 5G).

To examine whether PIEZO1-TMEM16F functional coupling directly regulates trophoblast fusion, we overexpressed PIEZO1 in TMEM16F KO BeWo cells and quantified forskolin-induced fusion. While PIEZO1 overexpression robustly enhanced fusion in WT BeWo cells (Fig. 4E-F), it failed to rescue the fusion deficit in TMEM16F KO cells (Fig. 5E-F). Our pressure clamp electrophysiology confirmed comparable PIEZO1 channel activity in WT and TMEM16F KO cells (Fig. S8), ruling out differential PIEZO1 expression or function as confounding factors. These results establish that PIEZO1’s pro-fusogenic role depends on TMEM16F, likely through TMEM16F-mediated PS exposure, a “fuse-me” signal required for trophoblast fusion ^5,23^. This functional coupling defines a mechanochemical axis where PIEZO1 senses mechanical cues and TMEM16F executes downstream biochemical signaling via PS externalization (Fig. 5G).

## Discussion

Our identification of PIEZO1 as a key regulator of trophoblast fusion expands its known functions in mechanobiology and development. Complementary to previous studies showing PIEZO1’s critical role in fetoplacental angiogenesis and embryonic development ^3,4,35–37^, our findings reveal that PIEZO1 in trophoblasts is also essential for embryonic survival. By manipulating PIEZO1 activity, our experiments demonstrate that mechanical forces play a pivotal role in driving trophoblast fusion during placental development.

Unlike endothelial PIEZO1, which responds to shear stress from blood flow ^38,39^, CTB-to-STB fusion mainly occurs at the basolateral side of STBs (Fig. 5G), distant from maternal and fetal circulation. This suggests that blood flow-induced shear forces are less likely to activate PIEZO1 at the fusogenic CTB-STB interface. Instead, trophoblasts may rely on local compressive forces at the fusogenic synapses ^40^, similar to those observed in myoblast fusion ^41,42^, to trigger PIEZO1 activation. Future investigation of these intrinsic mechanical cues will be crucial for understanding how PIEZO1 regulates trophoblast fusion.

Our *in vitro* experiments further demonstrate that PIEZO1-mediated Ca²⁺ influx activates TMEM16F, leading to surface exposure of PS, an essential ‘fuse-me’ signal required for trophoblast fusion ^5^. Notably, both PIEZO1 and TMEM16F are highly expressed in the SynT-2 layer (Fig. S2), and loss of either protein in trophoblasts severely disrupts the development of this STB layer, further supporting the importance of PIEZO1-TMEM16F coupling in trophoblast fusion (Fig. 5G). Beyond promoting TMEM16F-mediated PS externalization, PIEZO1-mediated Ca²⁺ influx may also activate additional signaling pathways that influence trophoblast gene expression, migration, and proliferation, processes that collectively impact trophoblast fusion, placental maturation, and overall embryonic development.

By identifying PIEZO1 as a crucial mechanosensitive ion channel in trophoblasts, our study lays a foundation for further exploration of trophoblast mechanobiology in placental physiology and pregnancy complications. Given the central role of trophoblast fusion in placental functions, dysregulation of PIEZO1 may contribute to pregnancy disorders such as preeclampsia, fetal growth restriction, preterm birth, and placental abruption. Future studies targeting PIEZO1 and its downstream pathways may offer therapeutic opportunities for managing these pregnancy-related complications.

### Materials and Methods Human placenta tissue

Human placental tissues were obtained under institutional review board approval (IRB# PRO00014627 and XHEC-C-2018-089), with a waiver of consent, to procure de-identified samples exclusively for research purposes. Placental specimens were fixed in 4% paraformaldehyde (Electron Microscopy Sciences, #15710) at 4°C for 48 hours. Following fixation, the tissues were preserved in 70% ethanol for 3 to 5 days and subsequently processed using a 4-hour protocol on the Leica ASP6025 tissue processor (Leica Biosystems). Processed tissues were paraffin-embedded using the HistoCore Arcadia system (Leica Biosystems) and sectioned at 5 μm thickness with a Leica RM2255 microtome (Leica Biosystems, Buffalo Grove, IL) for histological examination.

For analysis, tissue sections were de-paraffinized at 60°C, followed by two washes in xylene (Fisher Chemical, #X5S-4), and then rehydrated through a graded ethanol series (100%, 95%, and 70%). Sections were rinsed in MilliQ water for 5 minutes. Antigen retrieval was performed, after which tissue slices were permeabilized with 0.2% Triton X-100 and blocked with 10% goat serum prior to antibody incubation. PIEZO1 antibody (Proteintech, #82625-4-RR) was applied at a 1:200 dilution and incubated overnight. Fluorescent labeling was achieved using Alexa Fluor 488 fluorescence systems (Molecular Probes, #35552).

### Mice

PIEZO1-floxed ^27^ mice (B6.Cg-Piezo1^tm2.1Apat/J^, Strain #:029213) and Piezo1^P1-tdT^ ^4^ mice (B6;129-*Piezo1^tm1.1Apat^*/J, Strain #:029214) were obtained from the Jackson Laboratory or a kind gift from Bailong Xiao. To generate trophoblast-specific PIEZO1 knockout mice, *PIEZO1^flox/flox^*mice were crossed with *Elf5-Cre* mice ^28^. Genotyping was performed via PCR using DNA extracted from tail samples. Genomic DNA was extracted using an alkaline lysis method and PCR performed with GoTaq® Master Mix (Promega, #M7122). Amplified products were analyzed on a 2% agarose gel. All mouse handling and experimental procedures were conducted in strict accordance with the protocol approved by the Institutional Animal Care and Use Committee at Duke University (#A057-24-02) and complied with National Institutes of Health guidelines.

### Cell lines

The wildtype BeWo cell line was authenticated by the Duke University Cell Culture Facility and the TMEM16F knockout (KO) BeWo cell line was generated by sgRNAs targeting exon 2, as previously reported ^5,23^. BeWo cells were cultured in Dulbecco’s modified Eagle’s medium/Hams F12 (DMEM/F12) medium (Gibco, #11320-033) in a 5% CO_2_-95% air incubator at 37°C. The media were supplemented with 10% fetal bovine serum (FBS) (Sigma-Aldrich, #F2442) and 1% penicillin/streptomycin (Gibco, #15-140-122).

### siRNA transfection

BeWo cells were transfected with PIEZO1 SMARTPool siRNAs (Horizon, # M-020870-00-0005) using Lipofectamine RNAiMAX Transfection Reagent (Invitrogen, #13778075) following the manufacturer’s instructions. One day after siRNA transfection, the cells were switched to fresh medium and further cultured for another 24 hours before confocal imaging or fusion experiments.

### Fluorescence imaging of calcium and PS exposure

The procedures were done as previously described ^5,23^. Briefly, to detect calcium dynamics, BeWo cells were treated with 1 μM Calbryte 590 AM (AAT Bioquest, #20700) for 10 min at 37°C and 5% CO_2_. PS exposure was detected using 500 ng/ml CF 488-tagged AnV (Biotium, #29005). 10 µM Yoda1 (Cayman Chemical, #21904) and 5 µM GsMTx4 (MedChemExpress, #HY-P1410) were used to stimulate and inhibit PIEZO1 activity, respectively. Time-lapse imaging of BeWo cells was conducted before and after Yoda1 stimulation at room temperature using a Zeiss 780 inverted confocal microscope or an Olympus IX83 inverted epi-fluorescent microscope. ImageJ and a custom MATLAB code were used to quantify cytosolic calcium and AnV binding ^5^.

### BeWo cell fusion and quantification of fusion index

After BeWo cells were seeded on poly-L-lysine (Sigma-Aldrich)-coated coverslip for one day, cells were treated with 30 μM forskolin (Cell Signaling Technology, #3828s) for 48 hours to induce cell fusion. The forskolin-containing media was changed every 24 hours. After forskolin treatment, cells were stained with Hoechst (Invitrogen, #H3570, 1:2000) and Wheat Germ Agglutinin Alexa Fluor-488 Conjugate (Invitrogen, #W11261, 1:1000) for 15 min to visualize nuclei and cell membranes, respectively. Six random fields of view were acquired using a Zeiss 780 inverted confocal microscope for each group. Cell fusion was quantified by calculating the fusion index as previously described ^5,23^.

### PIEZO1 lentiviral overexpression

To overexpress PIEZO1, we utilize the lentiviral overexpression system as described previously ^43^. Briefly, we designed lentiviral vectors based on Addgene plasmid #62554 where the puromycin resistance gene was replaced with a hygromycin resistance gene from the pSBtet-GH plasmid (Addgene #60498) and the TMEM16F/Ano6 gene was replaced with the hPIEZO1 sequence (Genebank AGH27891.1) followed by an IRES sequence. For empty vector controls, hPiezo1-IRES was removed. All subcloning was done using an In-Fusion Snap Assembly (Takara #638947). Lentiviral constructs of PIEZO1 were packaged into lentivirus by transfecting HEK-293T cells with lentiviral vectors, pMD2.G, and psPAX2 (Addgene #12259 and #12260) using Lipofectamine 2000. WT and TMEM16F KO BeWo cells were then transduced with the filtered lentiviral supernatant and polybrene (10 µg/ml) and selected with hygromycin.

### Immunofluorescence staining

Cells were fixed with 1% PFA in phosphate-buffered saline (PBS) for 10 min, permeabilized with 0.1% Triton X-100 in PBS, and blocked with 5% goat serum in PBS for an hour. Coverslips were incubated in anti-PIEZO1 antibody (Proteintech, 15939-1-AP, 1:400) at 4°C overnight. A secondary antibody, Alexa Fluor 488 fluorescence system (Molecular Probes, #35552, 1:1000), was used for fluorescent staining. After staining nuclei with 4′,6-diamidino-2-phenylindole (DAPI), coverslips were mounted using ProLong Diamond Antifade Mountant (Invitrogen, #P36961) and imaged with a Zeiss 780 inverted confocal microscope.

### Mouse placenta histological analysis

Mice were deeply anesthetized using isoflurane. Placentas and embryos were freshly collected and fixed in 4% paraformaldehyde (Electron Microscopy Sciences, #15710) for 2 days at 4°C and then transferred to 70% alcohol for 3 to 5 days before being processed by using a 4-hour tissue processing setting with Leica ASP6025 (Leica Biosystems). Placentas were embedded in paraffin (HistoCore Arcadia H and Arcadia C, Leica Biosystems) immediately after the processing and then sectioned at 5 μm by using Leica RM2255 (Leica Biosystems, Buffalo Grove, IL) for histological staining.

H&E staining was performed through deparaffinization at 60°C followed by two washes with xylene (Fisher Chemical, #X5S-4) and a graded alcohol series (100%, 95%, 70%). The tissue was rinsed in MilliQ water for 5 min and then incubated in Hematoxylin 560 MX (Leica Biosystems, c#3801575) for 2 min, followed by rinsing with running water for 1 min. Tissue was dipped seven times in 0.3% ammonium hydroxide (Fisher Chemical, #A669S-500) and then rinsed in running water for 1 min. The tissue was incubated in 70% ethanol for 1 min followed by 10 dips in Alcoholic Eosin Y 515 (Leica Biosystems, #3801615). Dehydration was performed through an alcohol gradient of 95%, 100%, and twice in xylene, 1 min each. All slides were mounted using DPX Mounting Medium (Electron Microscopy Sciences, #13512). CD31 immunohistochemistry (IHC) staining was performed by using rabbit polyclonal to CD31 (Abcam, #ab28364) with the Avidin/Biotin Blocking Kit (Vector Laboratories Inc., #SP-2001), VECTASTAIN Elite ABC HRP Kit (Peroxidase, Standard) (Vector Laboratories Inc., #PK-6100), and VECTOR VIP Peroxidase (HRP) Substrate Kit (Vector Laboratories Inc., #SK-4600). Nuclei were counterstained with VECTOR Methyl Green (Vector Laboratories Inc., #H-3402-500). Microscopic imaging of histology samples was taken at 20X by using an Olympus microscope (BX63l, Olympus, Center Valley, PA) with cellSens Dimension software.

Immunofluorescence staining against MCT1 and MCT4 was performed on TMEM16F WT and KO placenta by using MCT1 antibody (Millipore-Sigma, #AB1286-I), which stains the fused SynT-1 layer facing the maternal blood sinuses, and MCT4 antibody (Santa Cruz Biotechnology Inc., #sc-376140), which stains the fused SynT-2 layer facing the fetal blood vessels. Immunofluorescence staining were imaged with a Zeiss 780 inverted confocal microscope

### Mouse placenta transcriptomic analysis

The reference clustering and individual single-cell RNAseq UMAP results were obtained from the Spatiaotemporal Transcriptomic Atlas of Mouse Placenta (STAMP, https://db.cngb.org/stomics/stamp/) ^26^.

### Electrophysiology

All ionic currents were recorded in the cell-attach configuration using an Axopatch 200B amplifier (Molecular Devices) and the pClamp software package (Molecular Devices). Glass pipettes were pulled from borosilicate capillaries (Sutter Instruments) and fire-polished using a microforge (Narishge) with a resistance of 2-3 MΩ for macroscopic current and 5-7 MΩ for single channel recordings. The bath solution consisted of the following (in mM): 140 CsCl, 10 HEPES, and 5 EGTA, with the pH adjusted to 7.4 using CsOH. For PIEZO1 experiments, the pipette solution contained (in mM): 140 CsCl, 10 HEPES, and 1 MgCl₂, with the pH adjusted to 7.4 using CsOH. PIEZO1 activation was achieved by applying pressure using a high-speed pressure clamp (ALA Scientific Instruments, #HSPC-2-SB). Pressure was applied using a step protocol starting from 0 mmHg to -60 mmHg with -10 mmHg increments. In the PIEZO1-TMEM16F coupling experiments, the pipette solution contained (in mM): 140 CsCl, 10 HEPES, 1 MgCl₂, and either 0 or 2.5 mM Ca²⁺, as indicated. The pH was adjusted to 7.4 using CsOH. The pressure was maintained at a constant value of -50 mmHg during the recordings to activate PIEZO1 and enable Ca^2+^ influx. A voltage step protocol from -100 mV to +160 mV in 20 mV increments was used to elicit TMEM16F channel activation. Cs⁺-based solutions were used to suppress contaminating K⁺ current in BeWo cells.

### Statistical analysis

All statistical analyses were performed using GraphPad Prism software. Unpaired two-tailed Student’s t-tests were used for single comparisons between two groups, while one-way ANOVA (with Tukey’s multiple comparisons test) was used for multiple comparisons. Sample number (n) values are provided in the figure legends. All data are presented as the mean ± standard error of the mean (s.e.m.).

## Acknowledgments

We sincerely appreciate the valuable insights and discussions provided by Drs. Carolyn Coyne and Liheng Yang. We thank Dr. Hua Pan for assistance with H&E and CD31 staining. We are also grateful to Drs. Haibin Wang and Bailong Xiao for generously providing the *Elf5-Cre* and Piezo1^P1-tdT^ mouse lines, respectively. We acknowledge the use of ChatGPT (https://chatgpt.com/) to assist with grammar checking and writing refinement in the Introduction and Discussion. Prompts used were [help check grammatical errors] or [analyze the following and suggest how it can be more concise without losing precision]. All final edits were reviewed manually to ensure accuracy and originality. BioRender was used to create the summary illustration.

## Author contributions

Conceptualization: HY, YZ

Methodology: YZ, KZS, PL, AJL, HY, LF

Investigation: YZ, KZS, PL

Visualization: YZ, KZS, PL, HY Funding acquisition: HY

Project administration: HY Supervision: HY

Writing – original draft: HY, KZS, PL, YZ

Writing – review & editing: all authors

## Data and materials availability

Original immunohistochemistry images are available at DOI: https://doi.org/10.7924/r4fj2qc4x. All remaining data are available in the main text or the supplementary materials.

