## Supplemental figures for "PIEZO1 Drives Trophoblast Fusion and Placental Development"

**Affiliations:**
[1]Department of Biochemistry, Duke University School of Medicine, NC 27710, USA
[2]Department of Obstetrics and Gynecology, Duke University School of Medicine, NC 27710, USA
[3]Department of Medicine, Duke University School of Medicine, NC 27710, USA
[4]Department of Neurobiology, Duke University School of Medicine, NC 27710, USA

† These authors contributed equally to this work
‡ Present address: Shenzhen Bay Laboratory, China

**This Supplementary file contains: 8 supplementary figures.**

### Supplementary Figures

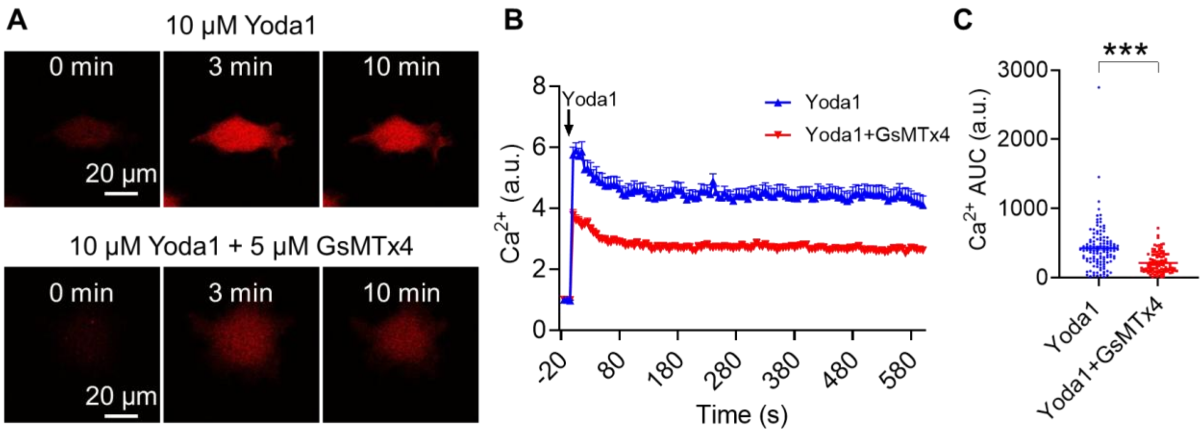


**Fig. S1. GsMTx4 inhibits Yoda1-induced $Ca^{2+}$ elevation in BeWo cells.**

**(A-B)** Representative images (A) and time course quantifications (B) of $Ca^{2+}$ increase in untreated control (top, n= 121) and 5 µM GsMTx4-treated BeWo cells (bottom, n=104) stimulated with 10 µM Yoda1. **(C)** Statistics of $Ca^{2+}$ elevation in (B). The area under the curve (AUC) was calculated from the $Ca^{2+}$ traces using GraphPad Prism, with the baseline set at $y$=1. *** $P<0.001$ (unpaired two-tailed t-test).

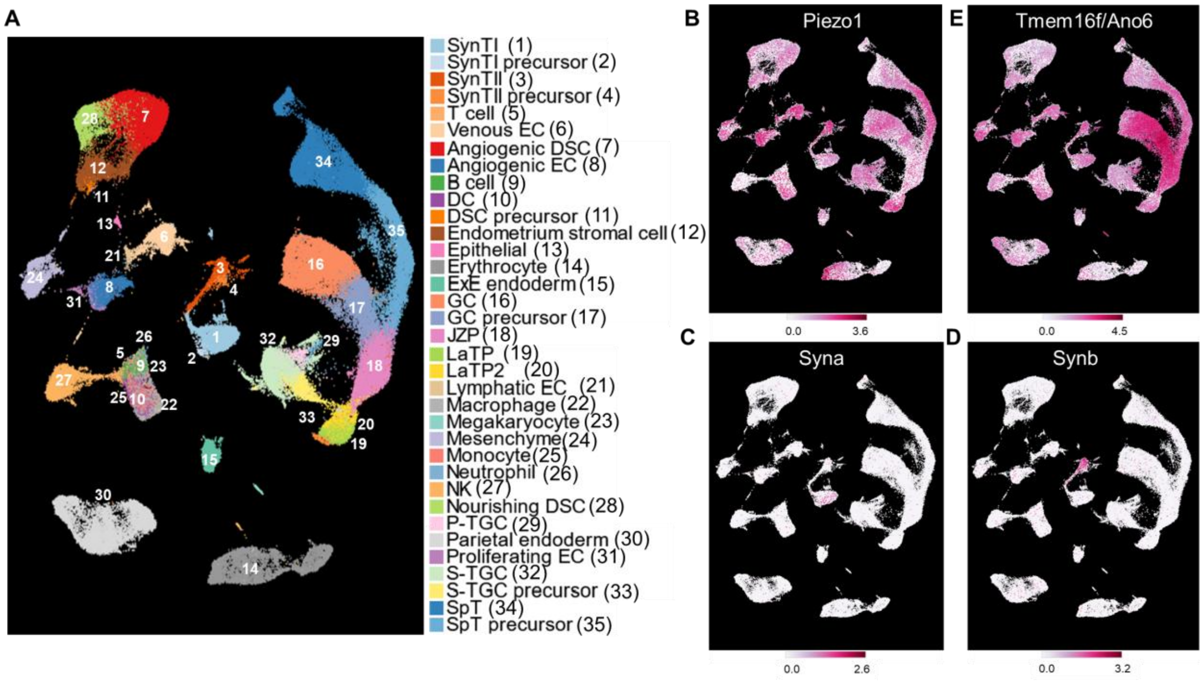


**Fig. S2. Transcriptomic expression of *Piezo1*, *Tmem16f*, *Syna* and *Synb* in the mouse placenta.**

**(A)** Reference UMAP plot of mouse placenta. **(B-E)** Single cell RNAseq UMAP plots of *Piezo1* (B), *Syna* (C), *Synb* (D), and *Tmem16f* (E). Data were extracted from the Spatiaotemporal Transcriptomic Atlas of Mouse Placenta (STAMP) [1].

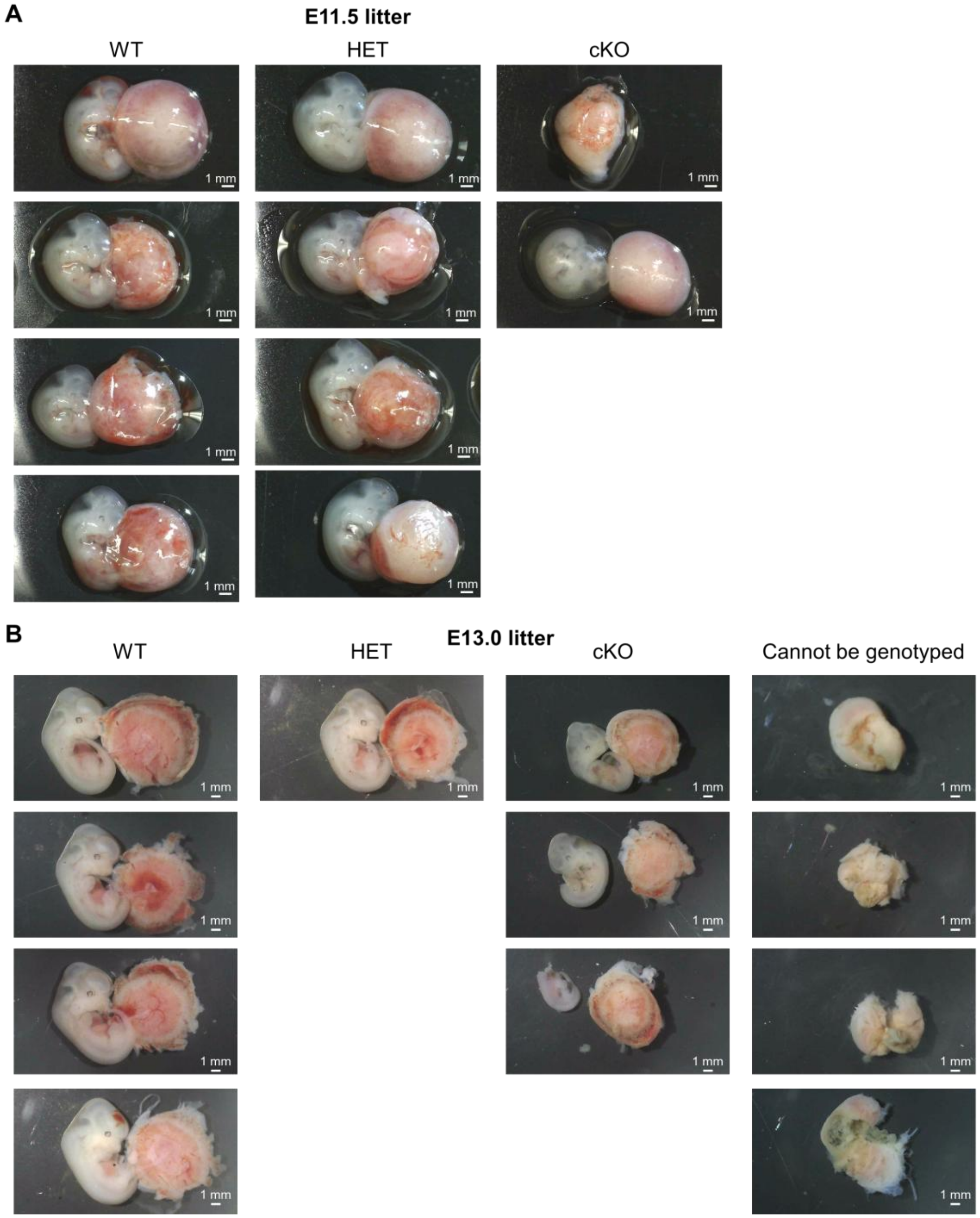


**Fig. S3. Morphology of Piezo1 conditional KO (cKO) placentas.**

**(A)** Representative images of an entire litter at E11.5. **(B)** Representative images of an entire litter at E13.0. Note that some embryos and placentas exhibit severe degradation and resorption. Genotyping could not be performed on these embryos due to the inability to clearly dissect intact embryonic tissues.

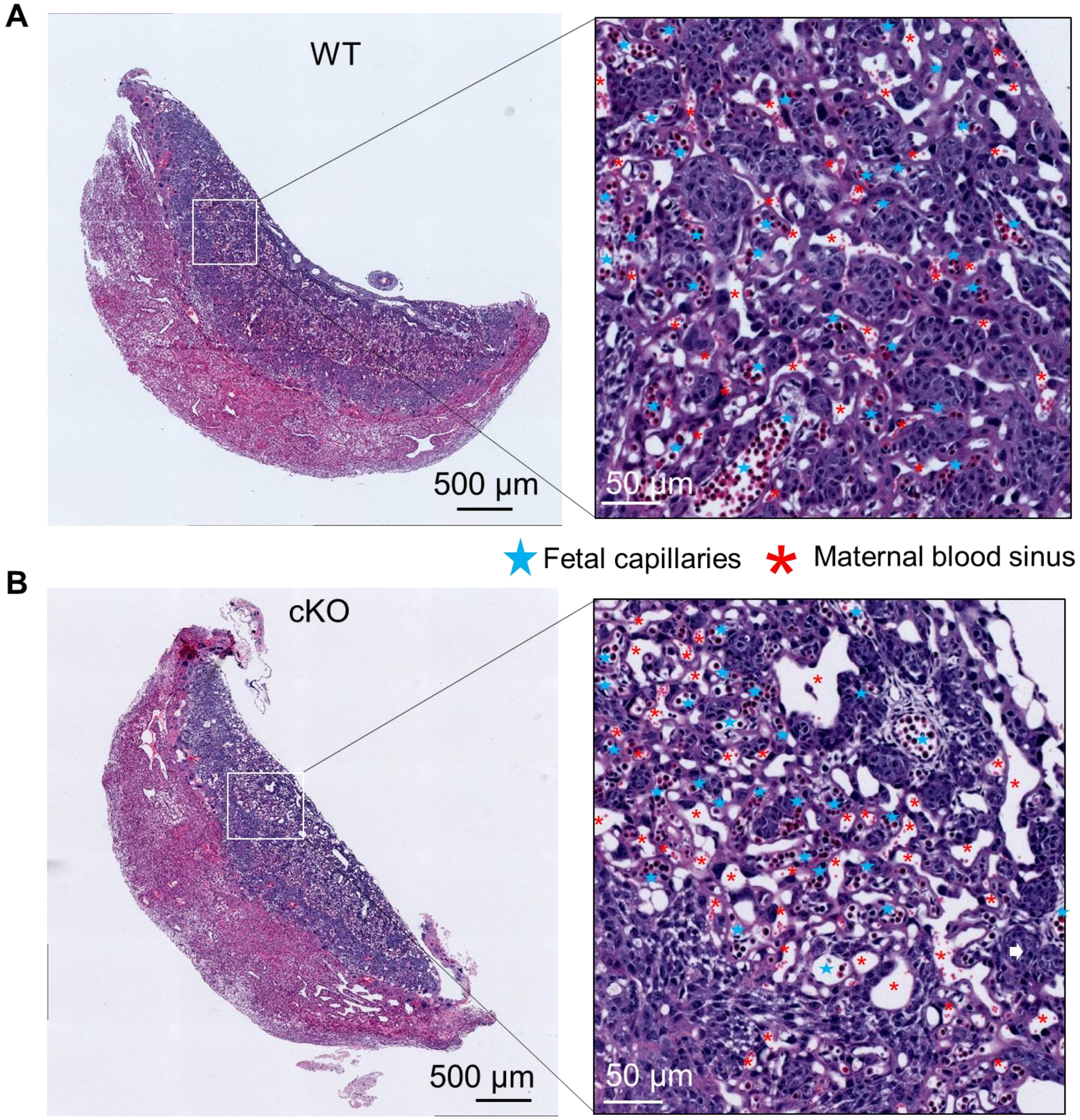


**Fig. S4. Mouse placenta H&E staining.**

**(A-B)** H&E staining PIEZO1 WT (A) and cKO (B) placentas at E13.0. The white boxes (left) were enlarged on the right. Red asterisks indicate maternal blood sinuses as evidenced by the presence of enucleated mature red blood cells. Blue stars denote fetal capillaries, characterized by the presence of nucleated fetal red blood cells. n = 5 biological replicates each for WT and cKO.

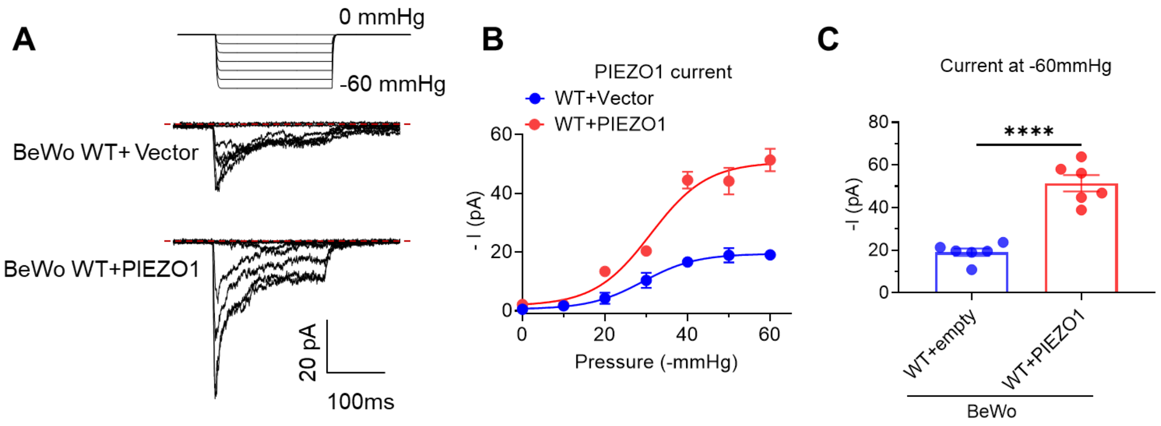


**Fig. S5. Validation of PIEZO1 overexpression in BeWo cells.**

5 **(A)** Representative cell-attached pressure-clamp recording of PIEZO1 currents from wildtype (WT) and hPIEZO1-overexpressed BeWo cells. The currents were elicited by pressure steps from 0 to −60 mmHg, with a holding potential set at −80 mV. **(B)** PIEZO1 current-pressure relationship on WT and hPIEZO1-overexpressed BeWo cells. Data are presented as mean ± s.e.m. n = 6 for both groups. **(C)** PIEZO1 current amplitudes at –60 mmHg from WT and hPIEZO1-overexpressed BeWo cells. Statistical analysis was done using an unpaired 2-sided Student’s t test. ****P < .0001. n = 6 for both groups.

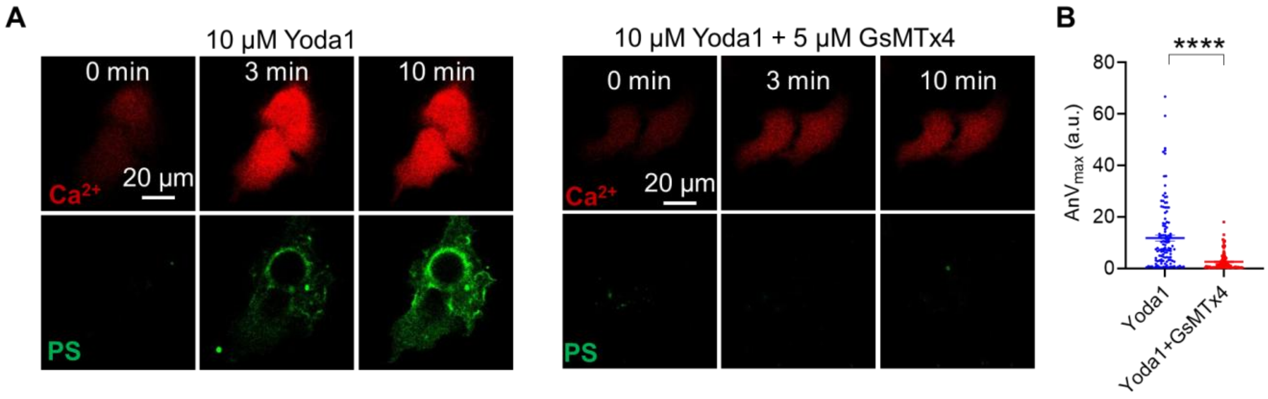


**Fig. S6. GsMTx4 inhibits PIEZO1-mediated PS exposure in BeWo cells.**

**(A)** Representative images of 10 μM Yoda1-induced $Ca^{2+}$ and PS exposure in untreated control (left) and 5 μM GsMTx4-treated BeWo (right). **(B)** Quantifications of the maximum fluorescence intensity of PS at 10 min post-Yoda1 stimulation for untreated control (n= 121) and GsMTx4-treated BeWo cells (n=104). Data are presented as mean ± s.e.m. Unpaired two-tailed t-test. ****P<0.0001.

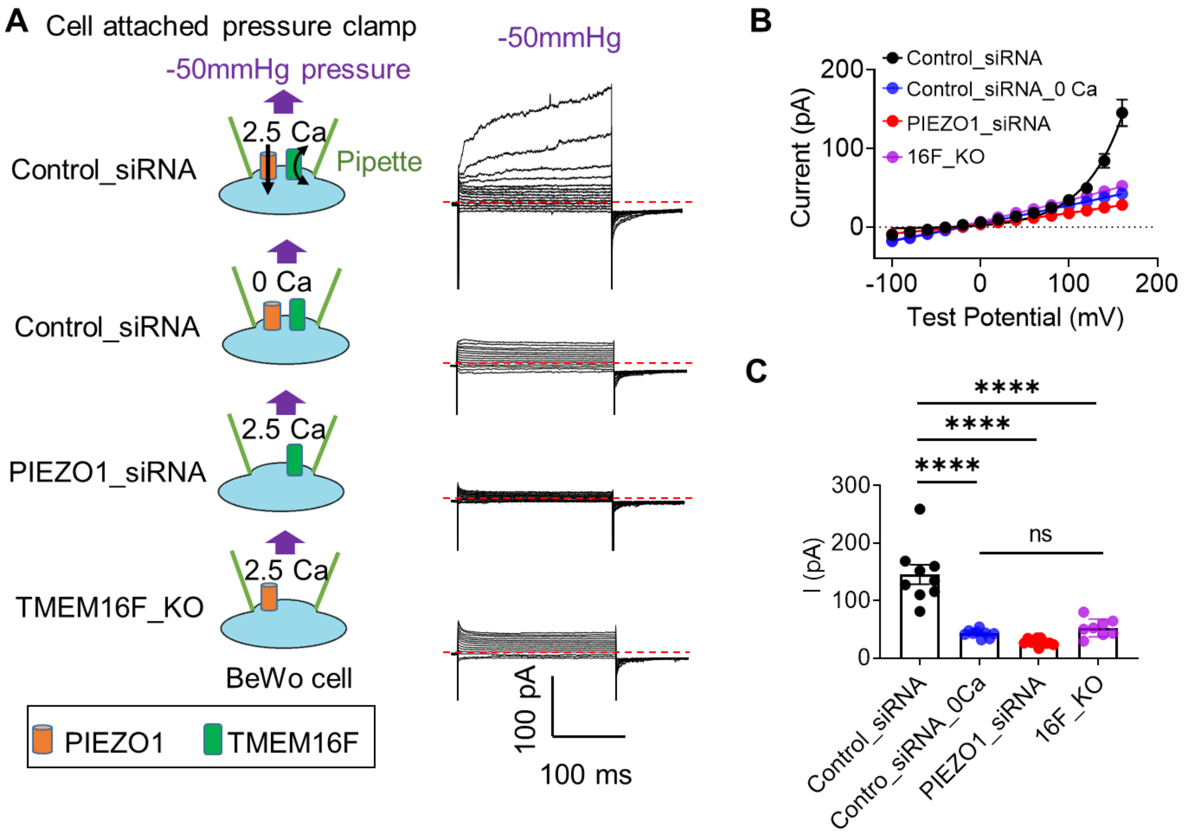


**Fig. S7. Mechanical stimulation of PIEZO1 activates TMEM16F.**

**(A)** Representative voltage clamp recordings under cell-attached mode were performed on BeWo cells under four conditions (from top to bottom): Control siRNA-treated WT cells (2.5 mM extracellular $Ca^{2+}$), Control siRNA-treated WT cells (0 extracellular $Ca^{2+}$), PIEZO1-siRNA treated WT cells (2.5 mM extracellular $Ca^{2+}$), and TMEM16F KO cells (2.5 mM extracellular $Ca^{2+}$). The current was elicited using a voltage step protocol from -100 mV to +160 mV under constant pressure of -50 mmHg. The schematics of the corresponding experimental conditions are displayed on the left. **(B)** I-V relationship of the pressure-induced current in (A) as indicated. **(C)** Comparison of the pressure-induced current +160 mV in (B). One-way ANOVA. The results are presented as mean ± s.e.m. **** $P < 0.0001$; n.s. not significant. n = 8-11 for each condition.

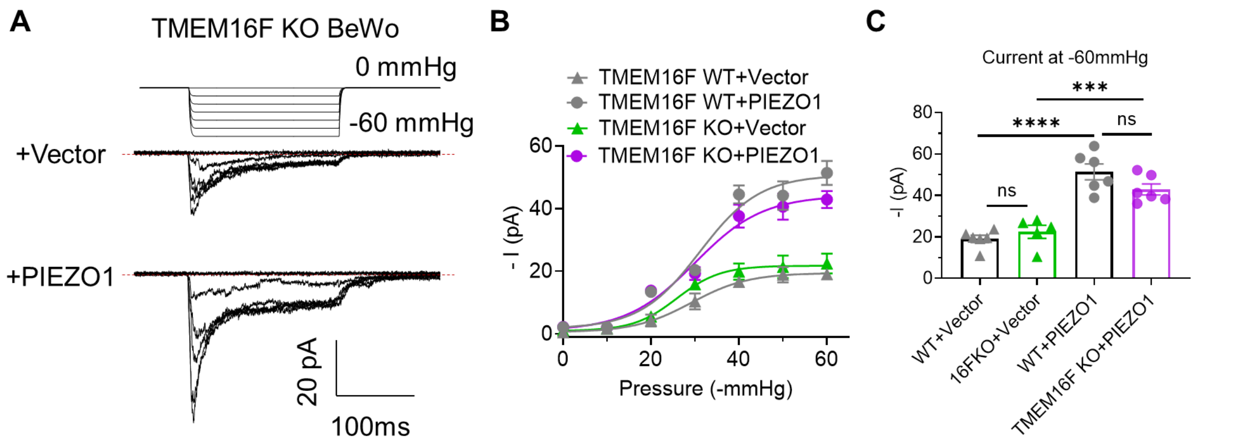


**Fig. S8. Quantification of PIEZO1 overexpression in TMEM16F KO BeWo cells.**

**(A)** Representative pressure-clamp recording of mechanosensitive current from TMEM16F-KO BeWo cells without (vector) and with hPIEZO1 overexpression. The current was elicited by a pressure clamp ranging from 0 to −60 mmHg, with a holding potential at −80 mV. **(B)** Current-pressure relationship of the currents recorded in (A). Mechanosensitive currents in WT BeWo cells with and without hPIEZO1 overexpression (gray symbols) were included for comparison. The results are presented as mean ± s.e.m. n = 5-6 for each condition. **(C)** Mechanosensitive current amplitudes at –60 mmHg from TMEM16F-WT and -KO BeWo cells without and with overexpression of hPIEZO1. Statistical analysis was done with one-way ANOVA. The results are presented as mean ± s.e.m. *** P < 0.001; **** P < 0.0001; n.s. not significant. n = 5-6 for each condition.
